# Z-scores unite >70 pairwise indices of ecological similarity and association for binary data

**DOI:** 10.1101/696047

**Authors:** Petr Keil

## Abstract

Pairwise ecological resemblance, which includes compositional similarity between sites (beta diversity), or associations between species (co-occurrence), can be measured by >70 indices. Classical examples for presence-absence data are Jaccard index or C-score. These can be expressed using contingency table matching components a, b, c and d - the joint presences, presences at only one site/species, and joint absences. Using simulations of point patterns for two species with known magnitude of association, I demonstrate that most of the indices describe this simulated association almost identically, as long as they are calculated as a Z-score, i.e. as deviation of the index from a null expectation. Further, I show that Z-scores estimated resemblance better than raw forms of the indices, particularly in the face of confounding effects of spatial scale and conspecific aggregation. Finally, I show that any single of the matching components, when expressed as Z-score, can be used as an index that performs as good as the classical indices; this also includes joint absences. All this simplifies selection of the “right” resemblance index, it underscores the advantage of expressing resemblance as deviation from a null expectation, and it revives the potential of joint absences as a meaningful ecological quantity.

## Introduction

Ecological resemblance (Legendre and Legendre 2012) is a fundamental ecological quantity that encompasses both similarity of species composition between sites (beta diversity) and co-occurrence patterns among species, which are the Q and R mode analyses respectively (Legendre and Legendre 2012, Arita 2017). Among the many measures of ecological resemblance, particularly popular are indices of pairwise similarity or association for binary presence/absence data, with around 80 proposed, as reviewed by (Hubálek 1982, Koleff et al. 2003, Rajagopalan and Robb 2005, Legendre and Legendre 2012, Ulrich and Gotelli 2013) (Appendix 1). These reviews provide guidance for the selection of appropriate indices, based on criteria such as symmetry, additivity, sensitivity to number of sites or species, or interpretability. Yet, the task of selecting the “right” index may still be daunting. Further, different researchers may deploy different indices, hindering synthesis and comparisons among studies.

All of the indices for binary data can be calculated using ‘abcd’ matching components, which follow contingency tables notation: a, the number of joint presences, b, the number of presences for only one species (or site), c, the number of presences for the other species (or site), and d, the number of joint absences. A classical example is Jaccard index (a/(a+b+c)) (Koleff et al. 2003) which has been used mostly in Q-mode analyses. Other examples are C-score (bc) and togetherness (ad) often used in R-mode (Ulrich and Gotelli 2013). However, all of the indices can be used in both Q- and R-mode, i.e. for rows or columns of a species-by-site incidence matrix (Hubálek 1982, Legendre and Legendre 2012, Arita 2017).

To standardize the indices, they can be expressed as *Z-score* (Gotelli and McCabe 2002, Ulrich et al. 2009, Ulrich and Gotelli 2013): *Z*=(*E*_*raw*_−*E*_exp_)/ *SD*_exp_, where *E*_raw_ is the index calculated on observed data, *E*_exp_ is a null expectation of the index, and *SD*_exp_ is standard deviation of the null expectation. This is similar to ‘standardized effect size’ in meta-analysis (Gurevitch et al. 1992, Ulrich et al. 2009), and it quantifies the deviation of the observations from a null expectation in common units of standard deviation. The null expectation from which *E*_exp_ and *SD*_exp_ are calculated is usually obtained from null models, i.e. by subjecting the data to a repeated randomization that breaks the association among species or sites. Currently, these null models are well developed particularly for binary species associations in R-mode (Gotelli 2000, Ulrich and Gotelli 2013), whereas for abundance data (Ulrich and Gotelli 2010) and for beta diversity assessments in Q-mode they are still being developed and their merits are unclear or debated (Chase et al. 2011, Ulrich et al. 2017, Legendre 2019).

While working on a related project, I noticed that several of the classical resemblance indices for binary data gave surprisingly similar answers when expressed as Z-scores, although in their raw form each of them might have a unique way of capturing resemblance (Koleff et al. 2003). This led me to a suspicion that that maybe the actual mathematical formula of resemblance indices does not matter that much, if at least some of the important matching components are in the formula, and as long as the index is expressed as a Z-score. To test this, and to figure out which of the components actually really need to be in an index, I devised a simulation exercise, as described below and in Appendix 1.

## Methods

I set this study to be in an R-mode, i.e. it examined the indices as measures of between-species associations (R-mode), and not beta diversity (Q-mode). The reason was that the null models are better developed for the R-mode [but see (Chase et al. 2011, Legendre 2019)] and it was more straightforward to simulate spatially explicit 2-species associations than patterns of beta diversity. However, I expect the same conclusions to be obtained for beta diversity (given the right null model is identified in the future), since the mathematics of the raw beta diversity is identical to the association metrics, only the indices are applied to a transposed site-by-species matrix (Arita 2017).

I simulated spatially explicit distributions of two species as two point patterns (Wiegand and Moloney 2014), while varying the magnitude of association between the two species, and while varying con-specific aggregation and number of individuals per species (detailed methods are in Appendix 2). I modelled the between-species association as being dependent on spatial distance, and it was controlled by a single parameter alpha, with negative values for segregation, zero for independence, and positive for attraction (Fig. S1, S2). I then aggregated the point patterns to grids of varying grain (resolution). My aim was then to recover this association alpha with various indices, or with their Z-scores, while expecting that varying grain, con-specific aggregation, and numbers of individuals to interfere with the ability of the indices to accurately recover that. Moreover, the limited ability of the indices to recover the undelaying association is also expected since the simulated association is realistically (Wagner 2003, Wiegand and Moloney 2014) distance-dependent, while the evaluated indices are naively spatially implicit.

I evaluated performance of 74 published indices of resemblance for binary data (Appendix 1) and of the individual four matching components abcd, all of them in their raw form and as Z-score. The performance was measured by Spearman rank correlation between the index and the true association alpha. I run one simulation for each combination of parameters and resemblance indices, resulting in 48,348 simulations in total. All code and data used are available at https://github.com/petrkeil/Z-scores.

## Results and Discussion

None of the indices gave perfect correlation with the association represented by parameter alpha (Fig. 1), which I expected since the association was distorted by varying conspecific aggregation, numbers of individuals, varying spatial grain, and the by fact that spatially implicit method was used for spatially explicit simulation. Expressing the raw indices as Z-score improved their median correlation with alpha from 0.72 to 0.78 (Fig. 1), and it dramatically reduced variation in the performance of the indices, from the interquartile range of 0.19 to 0.02 (Fig. 1). Thus, even though different indices give different answers concerning the magnitude of association between two species, they give very similar and good answers when used as Z-score.

**Figure 1.**
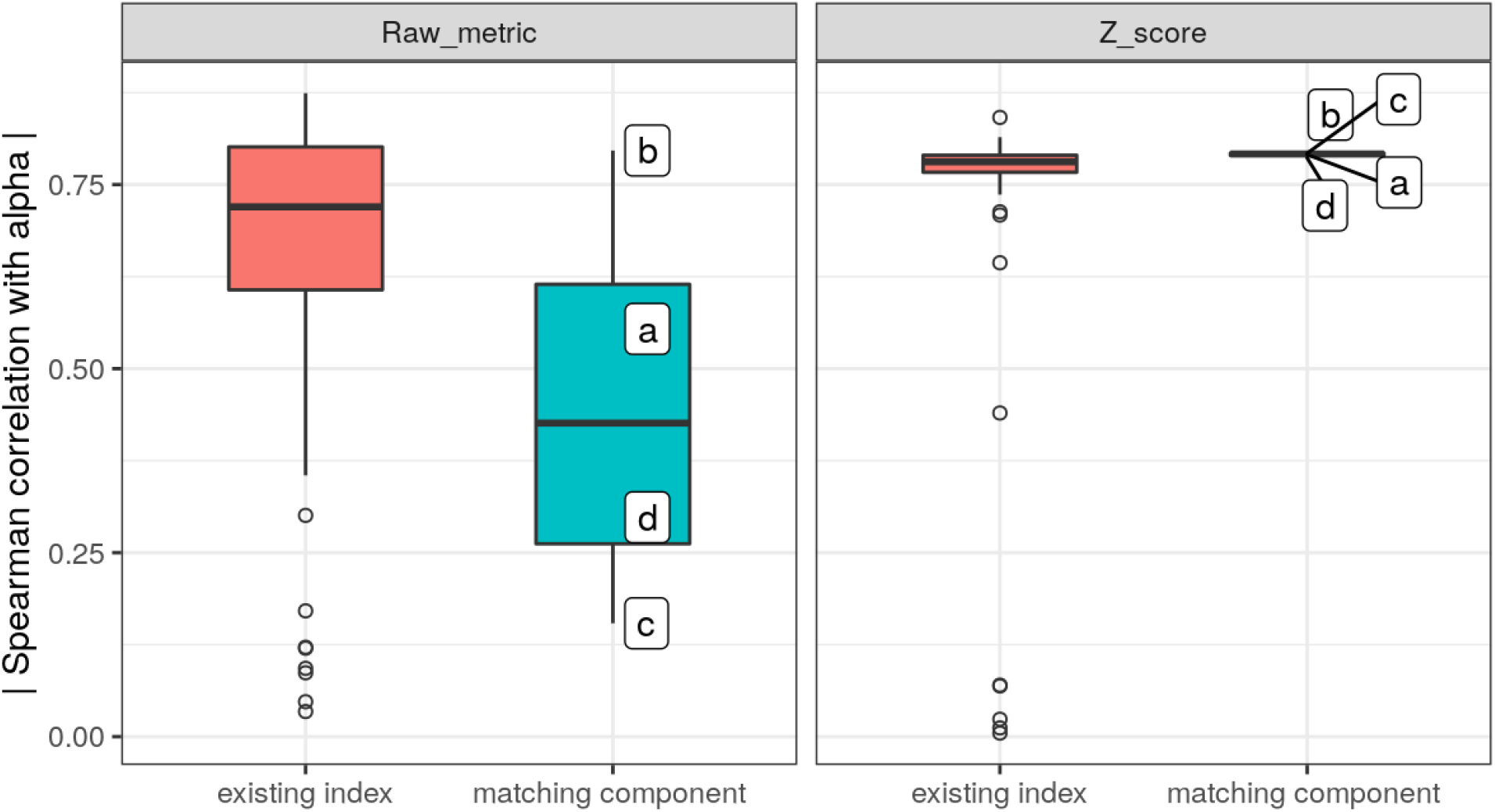
Performance of resemblance indices as measured by their Spearman correlation (its absolute value) with the parameter alpha that determines the magnitude of association (or segregation) of two simulated point patterns. Showed are the existing published indices for binary data (N = 74) and the four matching components a, b, c and d. Left panel shows raw values of the indices, right panel shows their Z-scores, i.e. deviation from a null expectation of no association.

The most striking result, however, was the performance of single matching components alone -- not only using them as Z-scores substantially improved their correlation with alpha (from median of 0.42 to 0.79), but they also gave almost identical answers (reduced interquartile range of the correlation from 0.35 to 0.003). Thus, any single one of the abcd components sufficiently captures the underlaying simulated association (Fig. 1). To explain this, in Appendix 3 I demonstrate that any single of the components is enough to capture the full magnitude of the actual species association. Surprisingly, this also includes the number of joint absences d (double zeroes), a quantity that has been deemed uninformative for ecological purposes (Legendre and Legendre 2012).

These results are reassuring to the field currently flooded with beta diversity and association indices. As long as an index is presented as a Z-score, it matters little which one is used - within the classical and most often used indices (Jaccard, Sorenson, Simpson, C-score), one can’t really go wrong, and one can even compare Z-scores from different studies, irrespectively of the actual index used, as long as the null model is the same and as long as it makes sense. Further, even indices that include double zeroes (e.g. the simple matching coefficient) are a good option, even though in their raw form they are heavily biased by any arbitrary addition of joint absences, e.g. due to arbitrary spatial delineation of the study area. I propose that this “united behavior” occurs because each of the matching components abcd carries equal information about the deviation from the null expectation (Fig. 1, Appendix 3). This leads me to a proposition that any of the matching components can be used as a perfectly valid index of ecological resemblance, if “Z-scored”, which has an alluring touch of minimalism.

I should stress that Z-scores indeed critically depend on the choice of the right null model. This is straightforward in the case of R-mode analysis of binary data, but less so in the case of Q-mode analysis (Chase et al. 2011, Ulrich et al. 2017, Legendre 2019), and it becomes a real problem when one has abundance data, due to the many ways such data can be randomized (Ulrich and Gotelli 2010). In a way, when a null model is deployed in order to move from the raw similarity index to the Z-score, one actually relaxes the need for the index to have a clear interpretation, since that shifts from the index to the null model. For example, Jaccard index is interpreted as proportional (or percentage) overlap of occurrences of two species, but it becomes a mere “summary statistic” (sensu Wiegand & Moloney 2014) when contrasted with expectations from a null model, where the null model now bears all the information/meaning. Moreover, in this paper I have demostrated that such an approach can actually improve our ability to capture meaningful association between two species, particularly in the face of confounding effects of spatial resolution of the data, uneven prevalences of the species, or intra-specific spatial aggregation. I thus argue that the null models and null expectations, perhaps more than particular indices, should be the central focus of future methodological research, particularly in the R-mode analyses of beta diversity.

## Acknowledgements

I am grateful to Jon Chase, Nick Gotelli, Arnošt Šizling, and David Storch for useful critical comments on an early draft of the manuscript. This work received support from German Centre for Integrative Biodiversity Research (iDiv) Halle-Jena-Leipzig funded by the German Research Foundation (FZT 118).

## Appendix 1 List of used indices

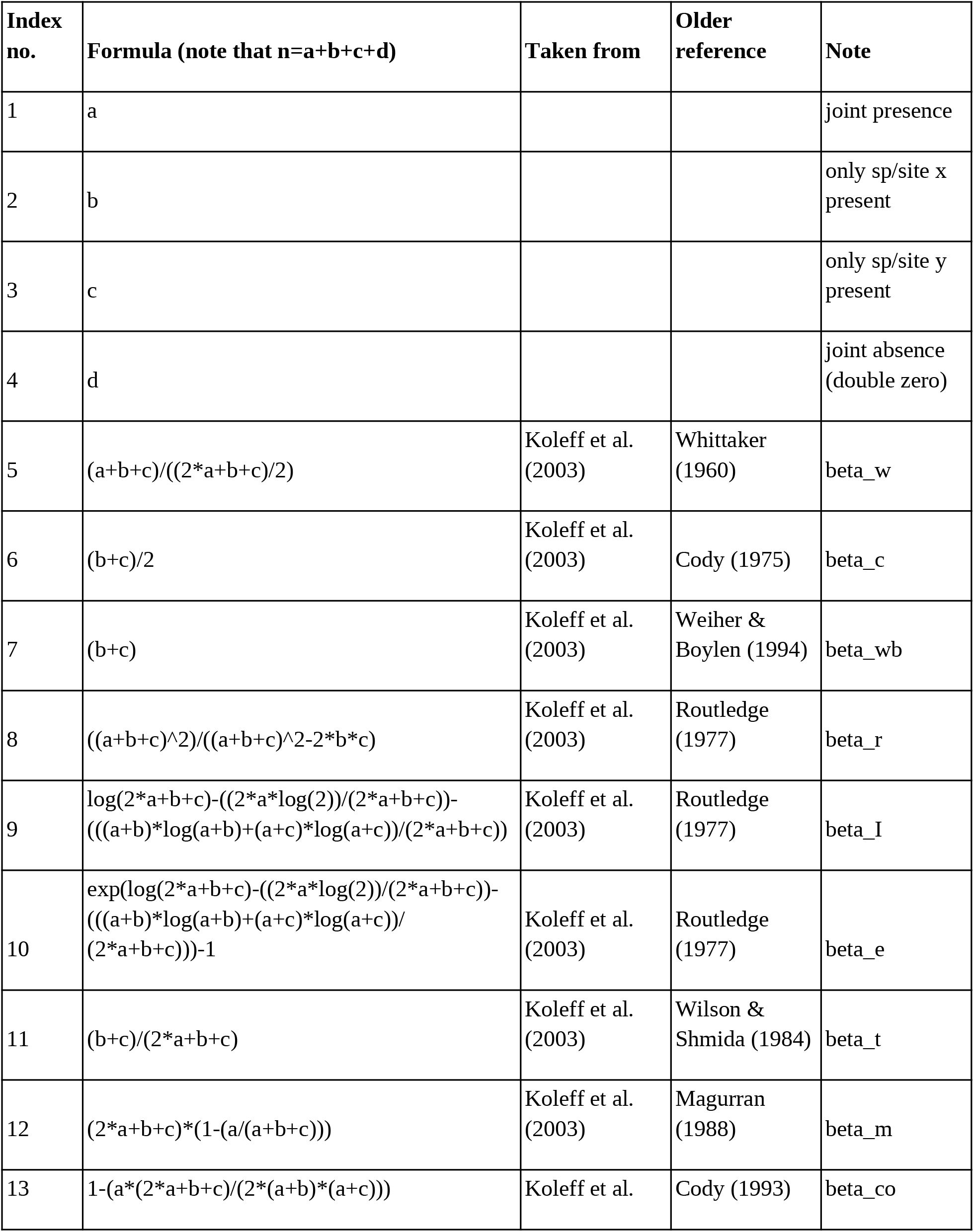
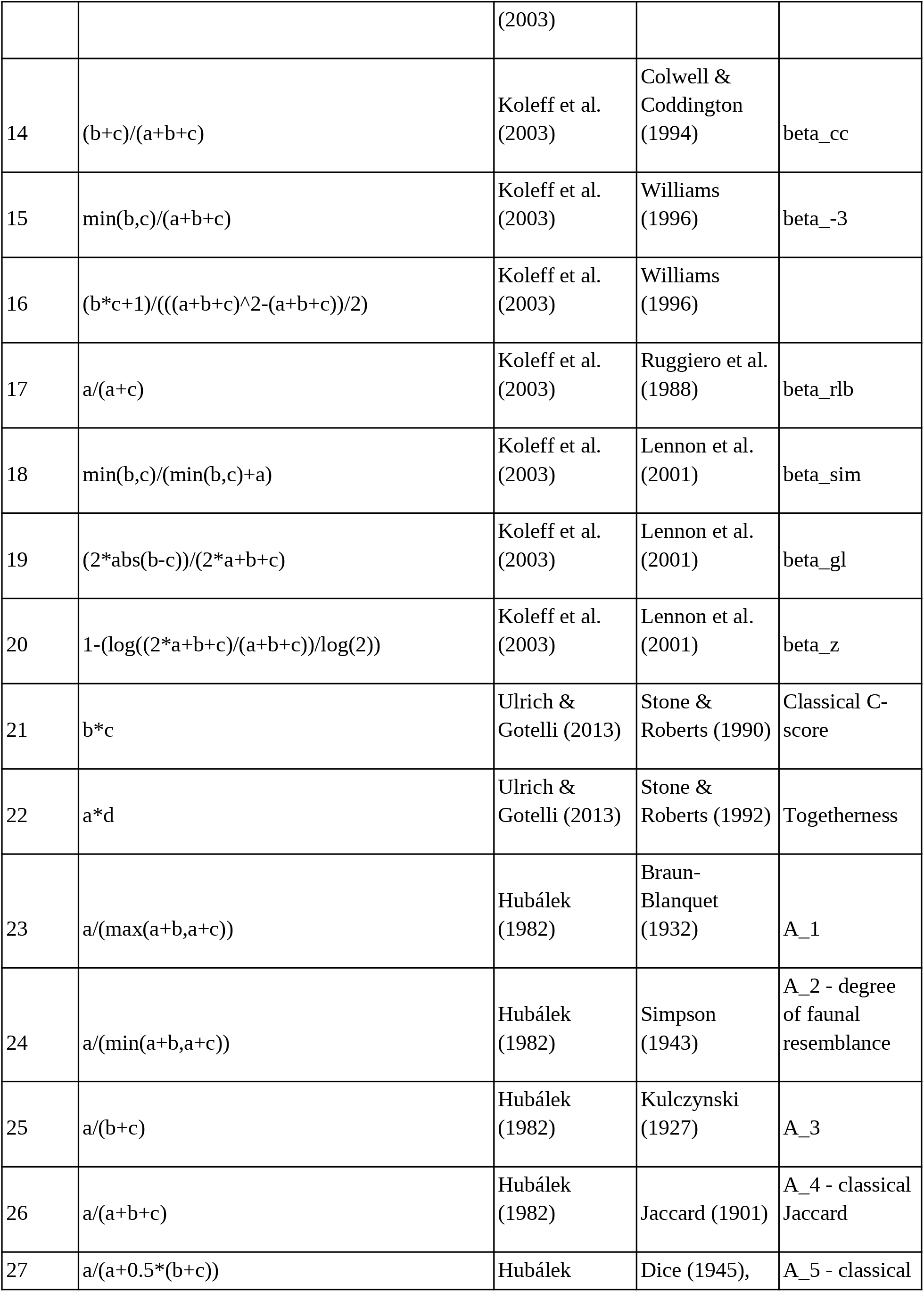
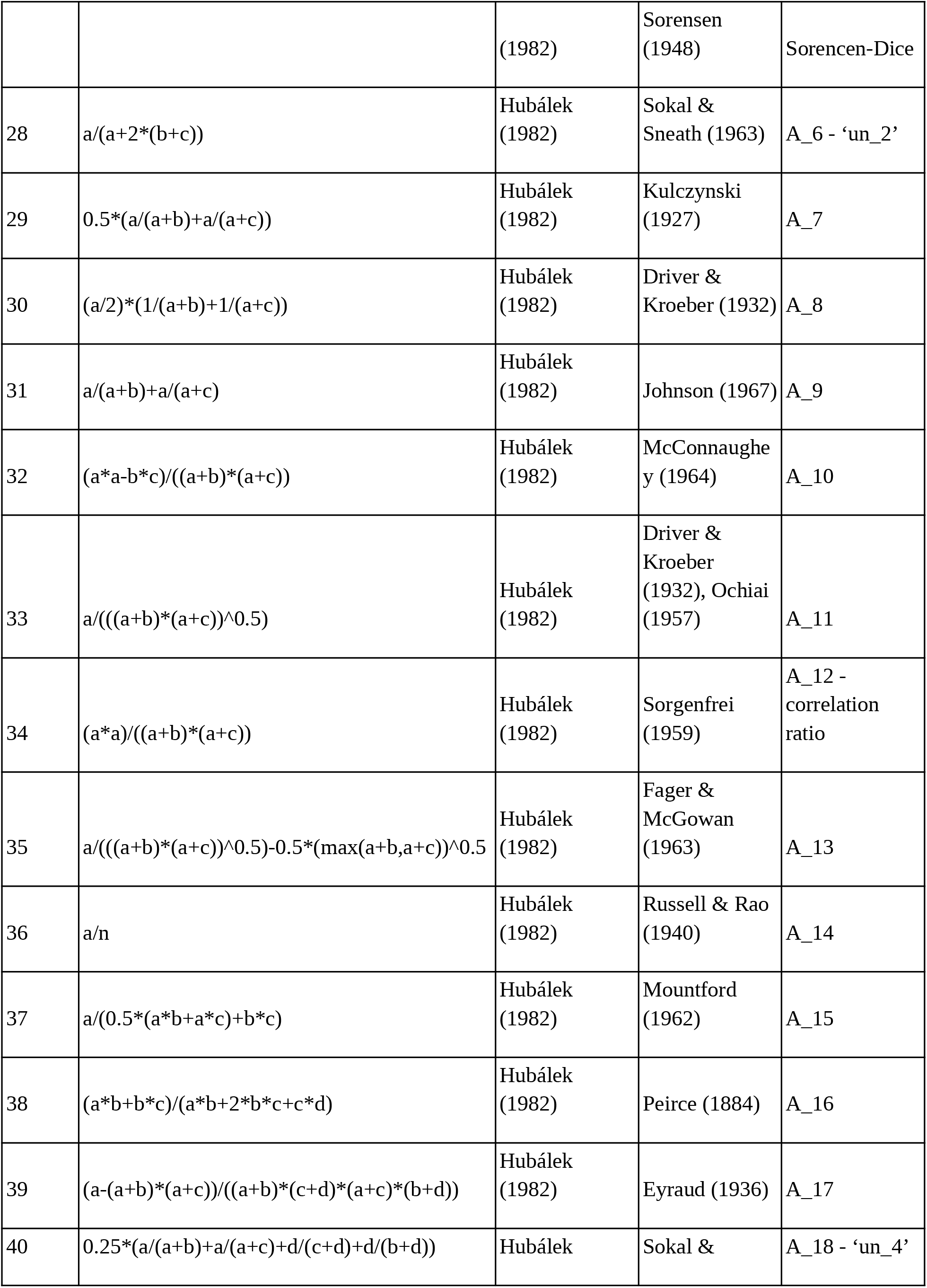
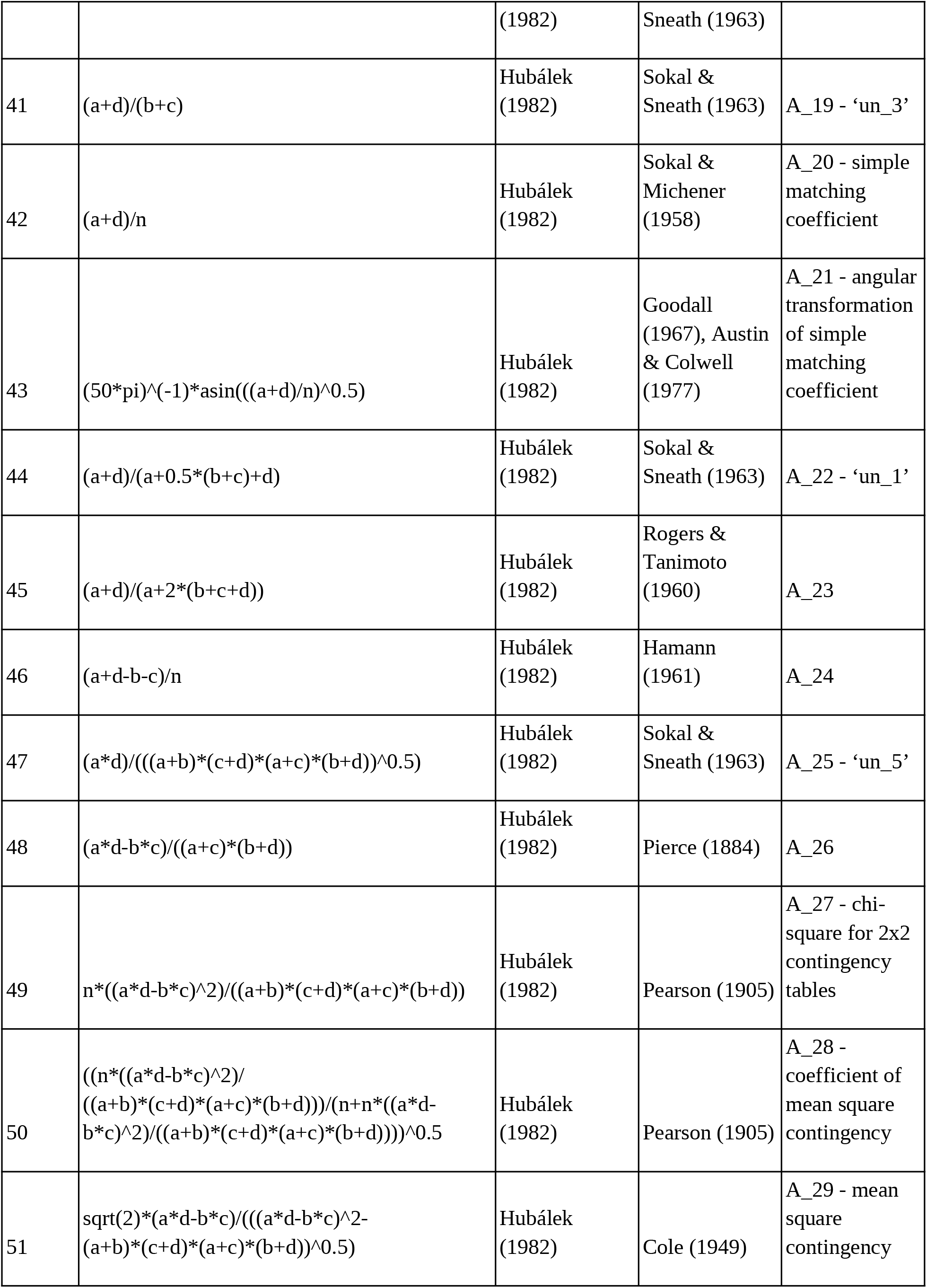
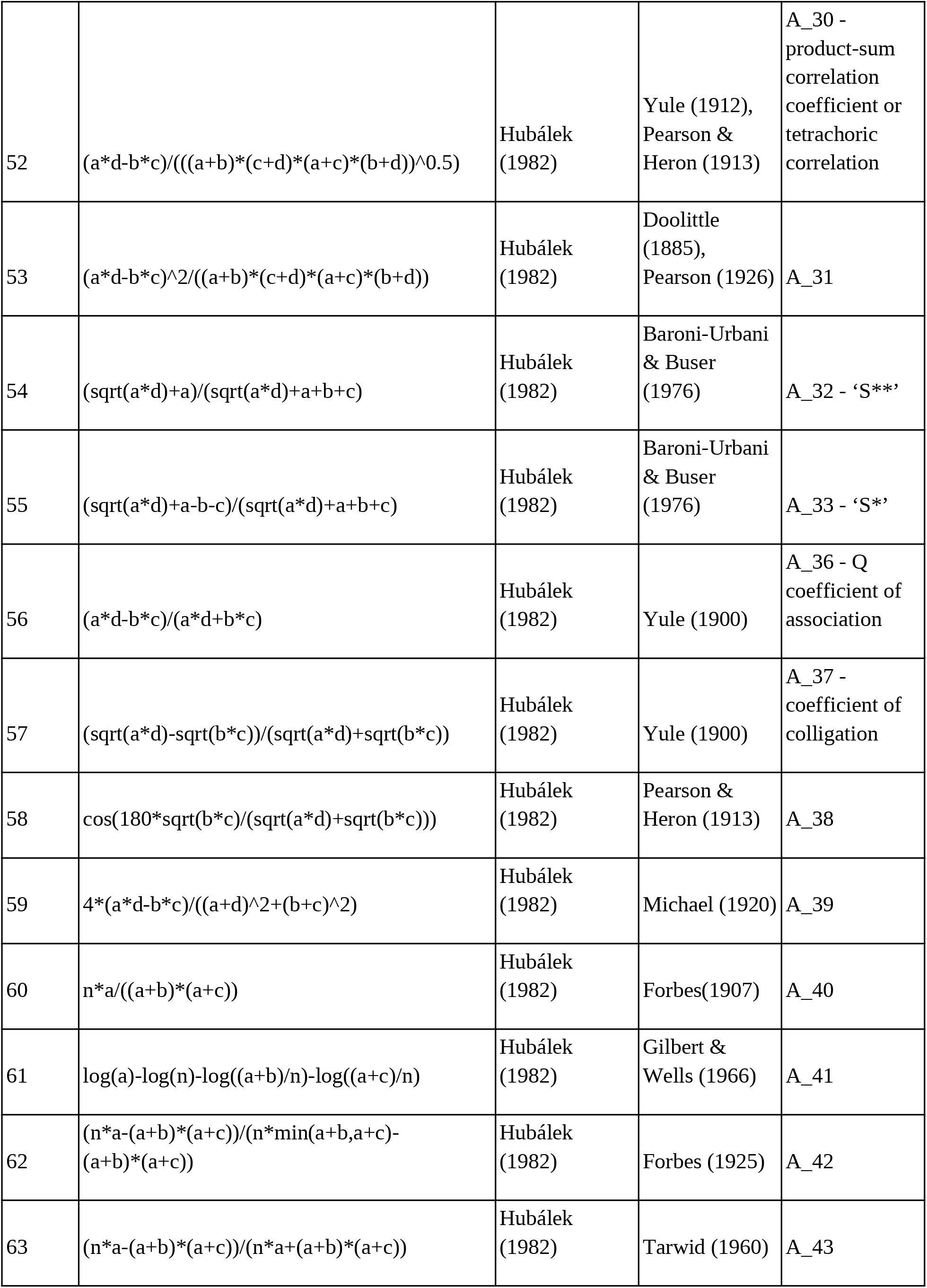
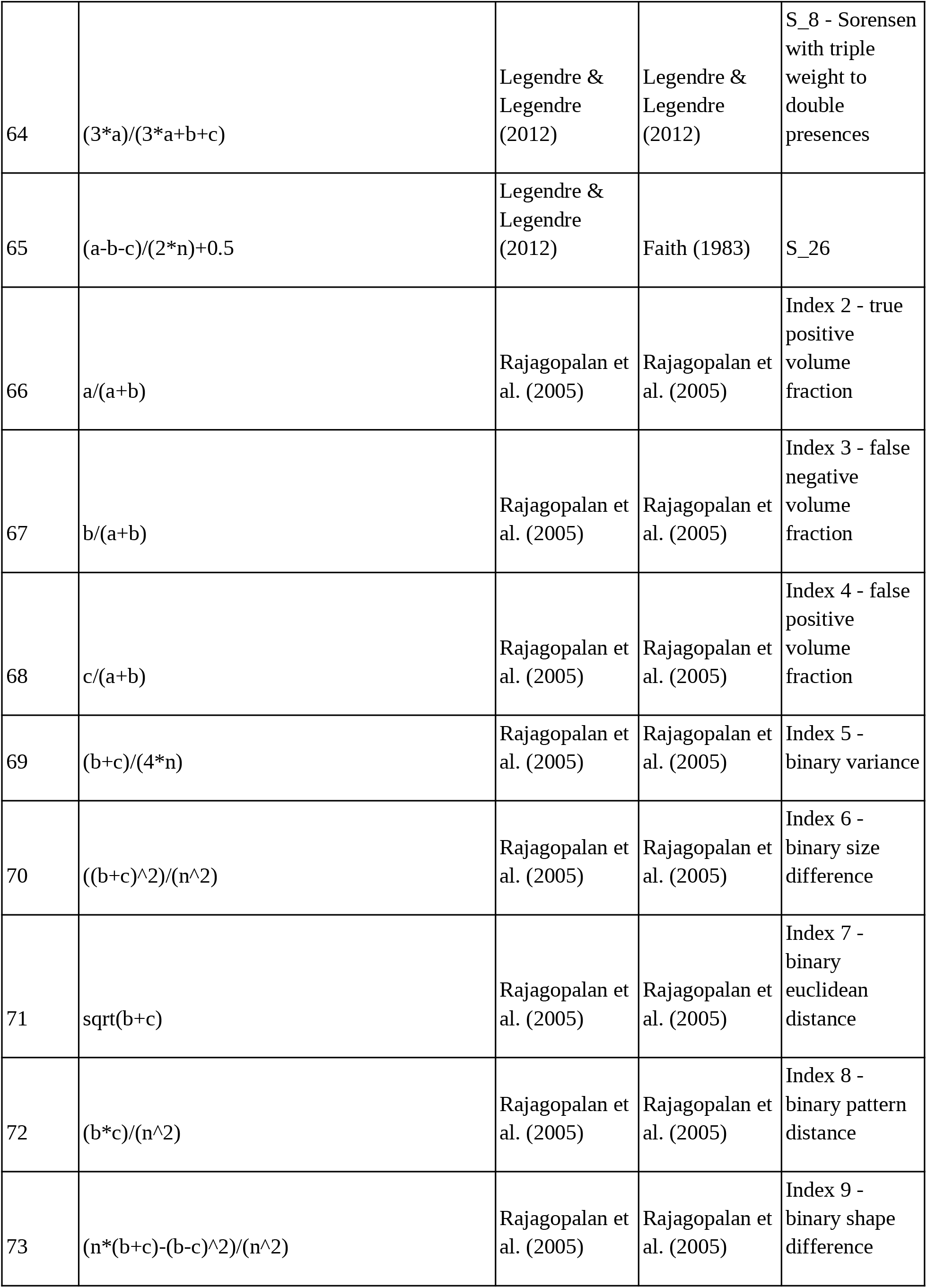
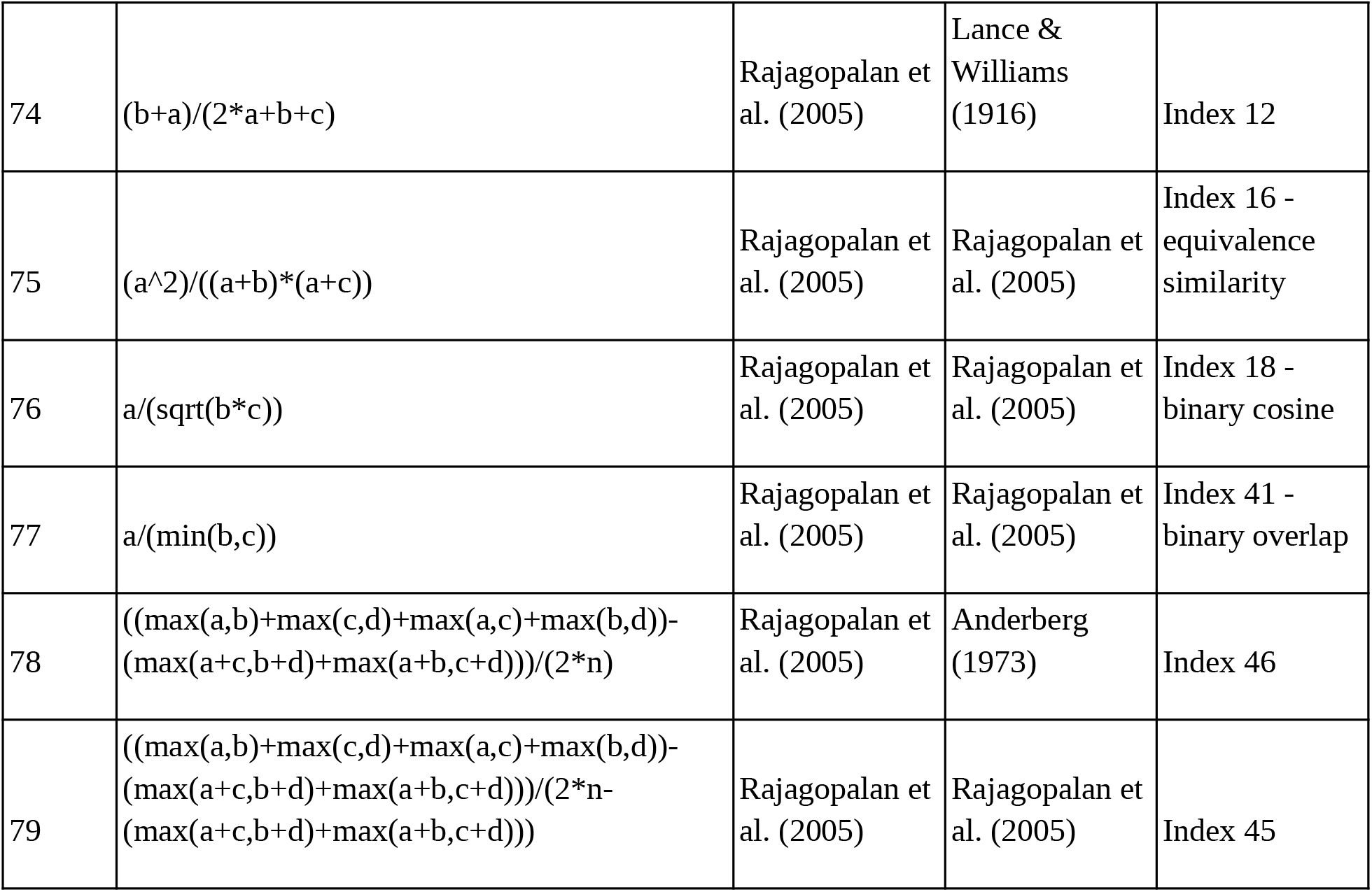

## Appendix 2 Detailed methods

### Exact simulation procedure

My goal was to devise simulations with known magnitude of inter-specific association, and to test how the different indices (or their Z-scores) are able to recover this association. Since the concept of association and co-occurrence is in reality inseparable from spatial distance, and thus any attempt to simulate realistic species distributions must be spatially explicit. I thus designed a simple simulation procedure where association between a species pair is controlled by a single parameter alpha, which can produce patterns of both segregation (alpha < 0), independence (alpha = 0), and attraction alpha > 0).

I simulated spatially explicit distributions of two species, sp1 and sp2 with abundances N1 and N2 respectively, as two point patterns in a square domain with side of 1. One simulation proceeded as follows (steps match Fig. S1):

(**a**) I chose a random point with coordinates*μ*_*x*_ and *μ*_*y*_ within the domain, with uniform probability density across the domain; this point was the center of distribution of sp1. (**b**) I created 2-dimensional probability density of points of sp1 as a bivariate normal distribution *f* _*sp*1_ (*μ,Σ*), where *Σ* is the covariance matrix with marginal variances *σ*_*x*_ = *σ*_*y*_ =*CSA*(con-specific aggregation) and with covariance *σ*_*xy*_ = 0. *μ* is the vector of coordinates *μ*_*x*_and *μ*_*y*_. (**c**) I drew N1 points from that probability density surface. (**d**) For every location in the domain I calculated its distance *r* from the nearest point of sp1. (**e**) I transformed *r* using a truncated exponential function (<u>Keil 2014</u>, Fig. S2b) 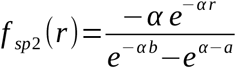 where *r* ∈ [*a, b*], where *a*=0 and *b*=1, but these truncation points can be set to any value depending on the size of the domain. The parameter *α* is the strength of interspecific association, with *α* <0 being segregation, *α* =0 independence, and *α* >0 positive attraction. (**f**) I drew N2 points from from the *f*_*sp*2_ (*r*). This whole procedure is visualized in Fig. S1, and examples of resulting patterns are in Fig. S2.

I simulated this model for each combination of the following parameter values:*CSA* ∈{0.001, 0.01, 0.1}, *N* 1 ∈{100, 1000}, *N* 2 ∈{100, 10000}, and for *α* ∈ {−20, −17.5, −15,…, 0,…, 15, 17.5, 20}, which I then aggregated to square spatial grids with {32, 16, 8} grid cells along each side.

### Z-score and null models

I expressed each of the indices (Appendix S2) as a Z-score (Ulrich et al. 2009, Ulrich and Gotelli 2013): *Z*=(*E*_*raw*_−*E*_exp_)/ *SD*_exp_, where *E*_*raw*_ raw is the index calculated on simulated data, *E*_exp_ is a null expectation of the index (below), and *SD*_exp_ is standard deviation of the null expectation. *E*_exp_ was calculated by subjecting the simulated raw co-occurrence data to a randomization procedure where incidences of within each species were reshuffled, assuming that all sites have equal probability of having an incidence; also, the total number of incidences in the reshuffled data is kept identical to the raw number of incidences. This is equivalent to the ‘sim2’ algorithm in R package EcoSimR (Gotelli et al. 2015). I repeat this procedure 400 times, and for each realization I calculated the index. *E*_exp_ was then the mean of the 400 realizations, and *SD*_exp_ was the standard deviation of these.

### Performance

Performance of each index, and its Z-score, was measured an absolute value of Spearman’s rank correlation. I used absolute value since some of the metrics are expressed as similarity, others as dissimilarity, but for my purpose this is irrelevant, and so taking an absolute makes the indices comparable.

**Figure S1.**
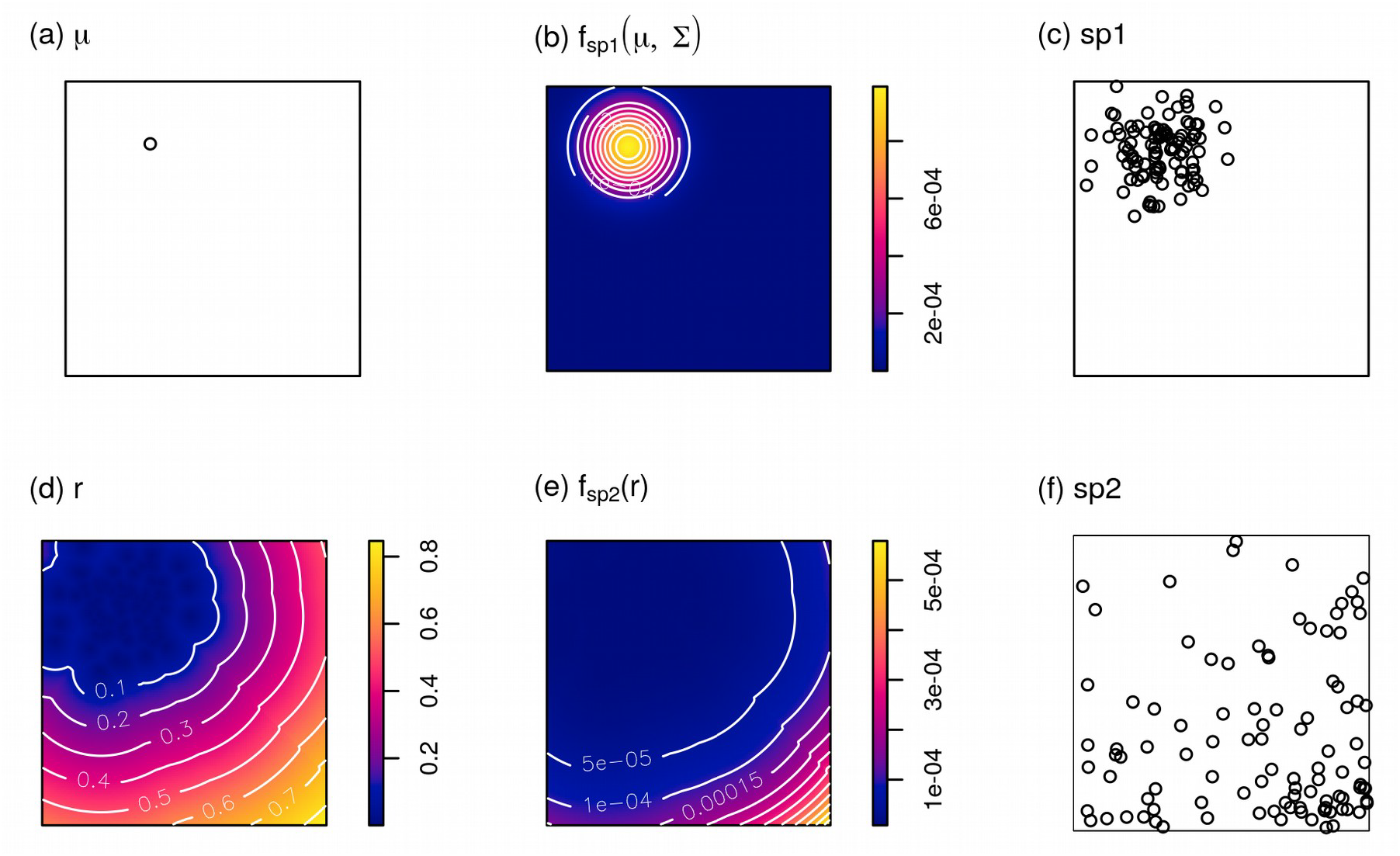
Steps of simulation of two point patterns of two species, sp1 and sp2, following the description in Appendix 1. Point pattern of sp1 is simulated as a point process with bivariate normal probability density *f*_*sp1*_ (*μ,Σ*) with zero covariance and marginal variances describing the con-specific aggregation (CSA) of sp1. Point pattern of sp2 is then simulated as a point process with *f*_*sp2*_ (*r*) describing the magnitude of inter-specific association alpha, where *r* is the distance to the nearest point of sp1.

**Figure S2.**
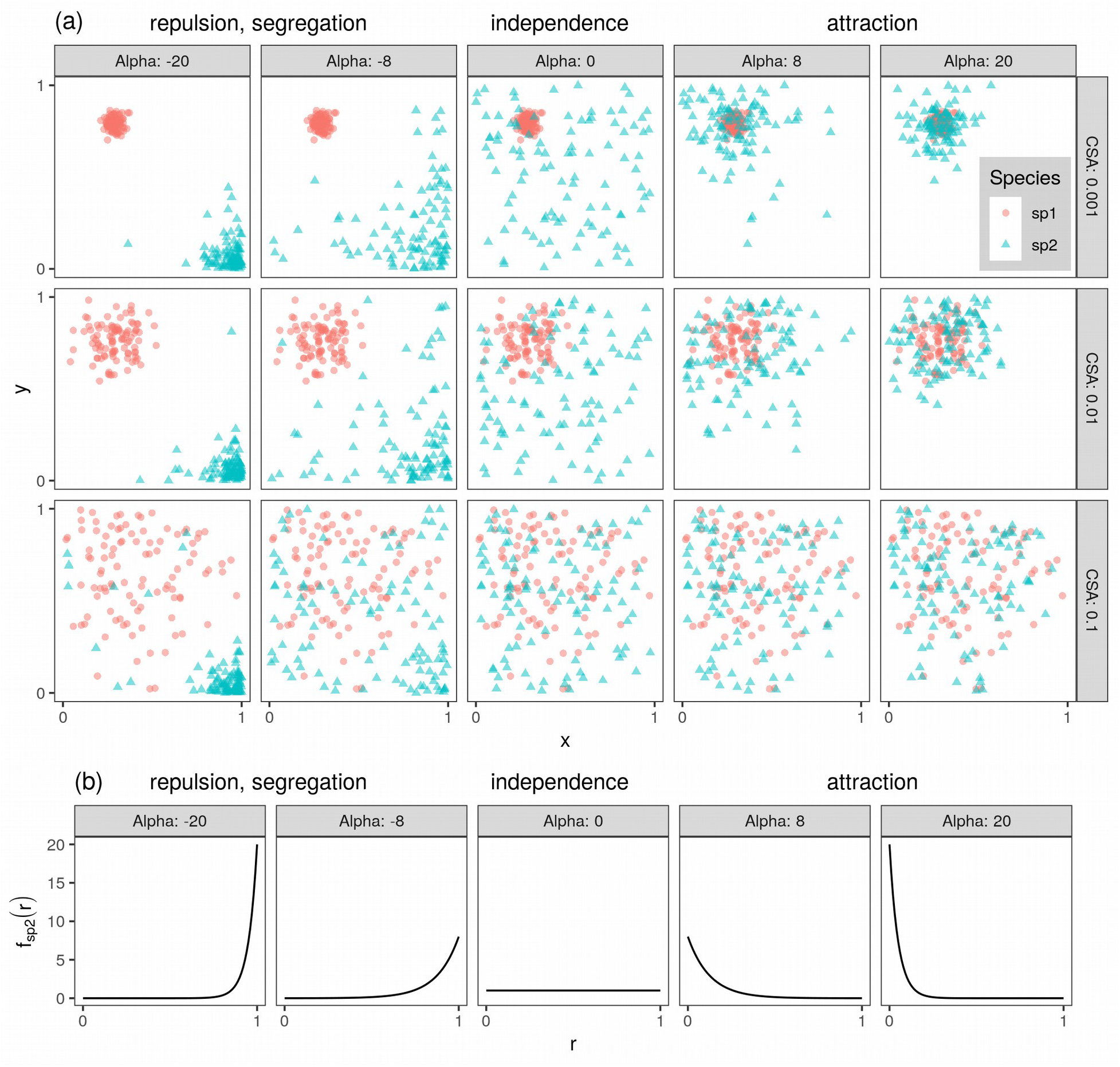
(a) Simulated spatial distributions of individuals (points) of two species (sp1 and sp2) in a square domain under 3 levels of con-specific aggregation of sp1 (CSA) and 5 levels of inter-specific aggregation *α* (alpha). (b) Truncated exponential probability density function (*f*_*sp2*_ (*r*), <u>Keil 2014</u>) describes how likely is to observe an individual of sp2 at a given distance *r* from any individual of sp1. This PDF is convenient since its shape depends on a single parameter (here called alpha) which represents various magnitudes of inter-specific repulsion (left) and attraction (right).

## Appendix 3 Demonstration that any single of the abcd matching components reflects true species association in the deviation of its observed value from null expectation

Let *x* and *y* be two species distributed over a set of sites. Let *a*_*obs*_ be the observed number of joint presences, *b*_*obs*_ the number of presences for only one species (or site), *c*_*obs*_ the number of presences for the other species (or site), and *d*_*obs*_ the observed number of joint absences. These can be arranged to a 2 *×*2 contingency table:

| | $x$ present | $x$ absent | |
| --- | --- | --- | --- |
| $y$ present | $a_{obs}$ | $c_{obs}$ | $n_y$ |
| $y$ absent | $b_{obs}$ | $d_{obs}$ | $a_y$ |
| | $n_x$ | $a_x$ | $N$ |

Where *n*_*x*_, *n*_*y*_,*a*_*x*_ and *a*_*y*_ are the marginal sums, and *N* is the grand sum of all counts in the table.

The expected abcd counts under a null expectation of independence with fixed row and column marginals are (Stevens 1938):

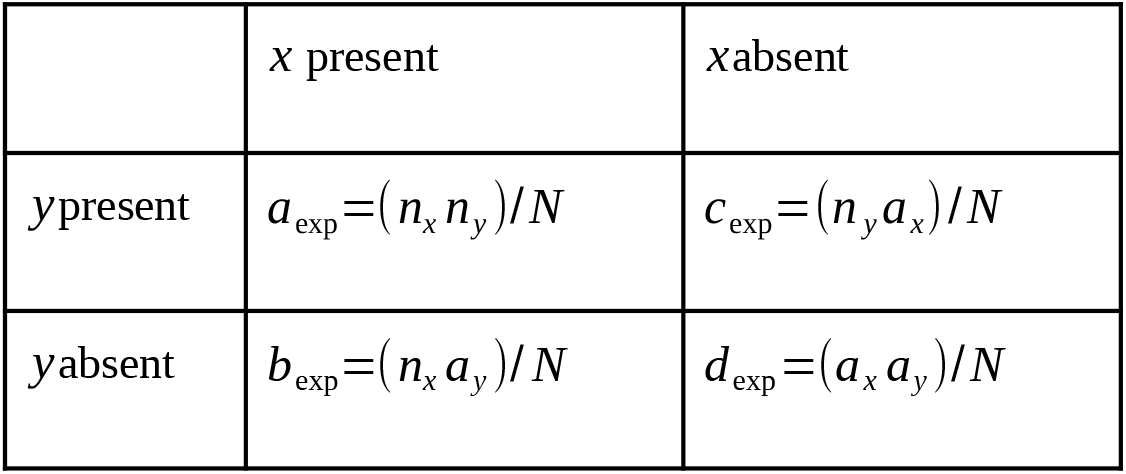

| | $x$ present | $x$ absent |
| --- | --- | --- |
| $y$ present | $a_{exp} = (n_x n_y) / N$ | $c_{exp} = (n_y a_x) / N$ |
| $y$ absent | $b_{exp} = (n_x a_y) / N$ | $d_{exp} = (a_x a_y) / N$ |

### Two species, two sites, occupancy of 1

Now consider two possible arrangements of the species by site binary incidence matrix with 2 sites and the two species *x* and *y*, where • indicates presence of a species, and empty cell is an absence. Each species occupies 1 site.

The first observed arrangement is *attraction*, where *a*_*obs*_=1, *b*_*obs*_=0, *c*_*obs*_=0, *d*_*obs*_=1

|  | Site 1 | Site 2 |
| --- | --- | --- |
| Species $x$ | ● | |
| Species $y$ | ● | |

The second observed arrangement is *segregation*, where *a*_*obs*_=0 *, b*_*obs*_=1 *, c*_*obs*_=1, *d*_*obs*_=0

The second observed arrangement is *segregation*, where $a_{obs}=0, b_{obs}=1, c_{obs}=1, d_{obs}=0$
|  | Site 1 | Site 2 |
| --- | --- | --- |
| Species $x$ | ● | |
| Species $y$ | | ● |

Using the (Stevens 1938) formulas above, the null expectation for both observed arrangements is *a*_exp_=*b*_exp_=*c*_exp_=*d*_exp_=0.5.

Thus any transition from aggregation to segregation will always lead to change in all four abcd matching components. Moreover, in both aggregation and segregation, all four observed components will always be different from the null expectation. Thus, any single of the four components can serve as an indicator of the overall magnitude of attraction or segregation in the matrix.

### Two species, four sites, occupancy of 2

Now let us move to a more complex case with species by site binary incidence matrix with 4 sites and the two species *x* and *y*, where • indicates presence of a species, and empty cell is an absence. Each species occupies 2 sites.

The first observed arrangement is *attraction*, where *a*_*obs*_=2 *, b*_*obs*_ =0, *c*_*obs*_=0, *d*_*obs*_=2

|  | Site 1 | Site 2 | Site 3 | Site 4 |
| --- | --- | --- | --- | --- |
| Species $x$ | ● | ● | | |
| Species $y$ | ● | ● | | |

The second observed arrangement is *partial overlap*, where *a*_*obs*_=1 *, b*_*obs*_=1, *c*_*obs*_=1, *d*_*obs*_=1

The second observed arrangement is *partial overlap*, where $a_{obs}=1, b_{obs}=1, c_{obs}=1, d_{obs}=1$
|  | Site 1 | Site 2 | Site 3 | Site 4 |
| --- | --- | --- | --- | --- |
| Species $x$ | • | • | | |
| Species $y$ | | • | • | |

The third observed arrangement is *segregation*, where *a*_*obs*_=0 *, b*_*obs*_=2 *, c*_*obs*_=2*, d*_*obs*_=0

The third observed arrangement is *segregation*, where $a_{obs}=0, b_{obs}=2, c_{obs}=2, d_{obs}=0$
|  | Site 1 | Site 2 | Site 3 | Site 4 |
| --- | --- | --- | --- | --- |
| Species $x$ | • | • | | |
| Species $y$ | | | • | • |

Using the (Stevens 1938) formulas above, the null expectation for all three of the observed arrangements is *a*_exp_=*b*_exp_=*c*_exp_=*d*_exp_=1.

We can see that the partial overlap is consistent with the null expectation of no association, which is intuitive.

Importantly, however, we can again see that *any* transition between any of the three arrangements leads to change of all four abcd components. For the Z-scores, what matters is the deviation from the null expectation, where again every single one of the four components reflects this departure.

### Conclusion

I have demonstrated that in both the 2-site and 4-site scenario, any deviation from the null expectation of independence must always be reflected by all four components abcd. I further dare to generalize the argument by stating that any observed site-by-species matrix consists of sub-matrices of the type described above (Ulrich and Gotelli 2013). Within these sub-matrices, any deviation from the null expectation will be reflected by all four abcd components. Thus, any of these four components can be used to quantify the deviation from the expected null expectation; what is even more useful that all four of them reflect both attractions and segregations. My simulations indeed corroborate this assertion -- as expected, all four components capture the true association very well, when expressed as a deviation from the null expectation.

